# Decoding Antimicrobial Resistance: Fusing Biologic Acumen and Computational Virtuosity for Paradigmatic Drug Innovation

**DOI:** 10.1101/2025.04.13.648583

**Authors:** Revanth Ananthoju, Parvatham Karthik Reddy

## Abstract

**Background:** Antimicrobial resistance (AMR) poses a critical threat to global health, necessitating innovative strategies to identify druggable targets and develop effective therapies. Traditional approaches often fail to address the rapid evolution of resistance mechanisms, particularly in high-priority pathogens. This study integrates AI-driven gene prioritization with nanotechnology to combat multidrug-resistant bacterial infections.

**Methods:** A dual-axis strategy was made to formulate genetic algorithm selection for gene centrality, conservation, interaction networks that has been effectively used to predict draggability scores for AMR genes (blaNDM-1, mcr-1). Hydroxyapatite-alginate-chitosan nano capsules were synthesized for pH-responsive ciprofloxacin delivery. Molecular docking validated ligand-target compatibility, while dynamic light scattering (DLS) and SEM characterized nanoparticle morphology (100–200 nm). In vitro drug release kinetics were assessed over 48 hours.

**Results:** blaNDM-1 showed an (draggability score: 0.92) and mcr-1 (0.89) that have been emerged as top targets with activity against both E. coli and Pseudomonas aeruginosa, due to their enzymatic roles in β-lactam and colistin resistance. Uniform nano capsules (150 ± 25 nm) demonstrated sustained ciprofloxacin release (75% cumulative release at 24 h). Docking confirmed stable binding of ciprofloxacin to active sites of target proteins (hydrogen bonds ≤2.0 Å), supporting inhibitory potential.

**Conclusions:** These findings indicate computational and experimental approaches to address AMR. The AI-nano therapy pipeline identified high-value targets and validated nanoparticle-based drug delivery, achieving a 6-log reduction in bacterial load in vitro. Future work will focus on in vivo validation and clinical translation.

## 1. Introduction

The rise of antimicrobial resistance (AMR) in Pseudomonas aeruginosa and Escherichia coli has severely limited therapeutic options for infections caused by these multidrug-resistant (MDR) pathogens. Carbapenems, once considered last-line agents against Gram-negative bacteria, are increasingly compromised by resistance mechanisms such as metallo-β-lactamase (MBL) production (e.g., blaNDM-1) and efflux pump overexpression [1, 2]. P. aeruginosa, in particular, exploits biofilm formation and quorum sensing (QS) systems to evade antibiotics, further complicating treatment [3]. Biofilms, encased in extracellular polymeric substances (EPS), act as physical barriers and promote persistent infections, while QS-regulated virulence factors like elastase (lasA/lasB) and rhamnolipids (rhlA) enhance pathogenicity [4, 5].

Ciprofloxacin, a broad-spectrum fluoroquinolone, remains a critical agent against P. aeruginosa and E. coli.

However, its efficacy is undermined by resistance mechanisms such as DNA gyrase mutations and efflux pump activation, as well as systemic toxicity and poor biofilm penetration [6, 7]. To address these limitations, nanotechnology has emerged as a promising strategy to enhance drug delivery. Nanoparticles can improve bioavailability, target biofilm microenvironments, and reduce off-target effects by leveraging stimuli-responsive release mechanisms [8]. Despite these advances, conventional approaches often lack precision in identifying high-value AMR targets and optimizing delivery systems for resistant pathogens. This study introduces an innovative dual-axis strategy that synergizes artificial intelligence (AI)-driven target prioritization with nanotechnology-enhanced drug delivery. resistance occurred due to changes in porin expression, rendering the outer bacterial membrane impermeable to imipenem [2]. First, a genetic algorithm analyzed gene centrality, conservation, and pathway relevance using data from the Comprehensive Antibiotic Resistance Database (CARD) and Drug Bank to predict druggable AMR targets. This approach identified blaNDM-1 (a Carbapenemases gene in P. aeruginosa) and mcr-1 (a colistin resistance gene in E. coli) as top candidates due to their enzymatic roles and clinical prevalence.

Second, pH-responsive hydroxyapatite-alginate-chitosan nano capsules were engineered to deliver ciprofloxacin. These biocompatible carriers exploit the acidic biofilm microenvironment to trigger drug release, enhancing penetration and retention while minimizing systemic toxicity. Molecular docking validated ciprofloxacin’s stable binding to target proteins (hydrogen bonds ≤2.0 Å), confirming its compatibility with prioritized AMR enzymes. Dynamic light scattering (DLS) and scanning electron microscopy (SEM) characterized the nanoparticles, revealing uniform morphology (150 ± 25 nm) and sustained drug release (75% cumulative release at 24 hours). In vitro assays demonstrated a 6-log reduction in bacterial load for both P. aeruginosa and E. coli biofilms, underscoring the formulation’s potency. By integrating computational precision with nanoscale delivery, this work bridges critical gaps in AMR therapeutics, offering a scalable framework to combat infections caused by these priority pathogens. This research not only advances targeted drug discovery but also redefines the role of nanotechnology in overcoming biofilm-mediated resistance. Future studies will focus on in vivo validation and clinical translation to address the growing global burden of AMR.

## 2. Material and methods

### 2.1. Ethics approval and informed consent

This study utilized publicly available genomic data from the Comprehensive Antibiotic Resistance Database (CARD) and Drug Bank, adhering to ethical guidelines for computational research. Clinical isolates of Escherichia coli (ATCC 25922) and Pseudomonas aeruginosa (ATCC 27853) were obtained from the American Type Culture Collection (ATCC). No human or animal trials were conducted, ensuring compliance with institutional ethical standards.

### 2.2. Materials

All chemicals and reagents employed for nanoparticle synthesis were commercially available and conform to the standards used in similar investigations. For example, key components such as hydroxyapatite and chitosan (with a molecular weight of approximately 50 kDa) were sourced from well-established chemical suppliers, and sodium alginate along with calcium chloride were acquired from reputable vendors. Ciprofloxacin, the antimicrobial agent chosen for this study, was similarly procured from a commercial source. All solvents and buffers utilized in the experiments were of analytical grade, ensuring high purity and consistency.

### 2.3. Instruments and Computational Tools

The physicochemical characterization of synthesized nanoparticles included a Dynamic Light Scattering (DLS) analyzer and a Scanning Electron Microscope (SEM), which are standard tools in nanoparticle research. Computational analyses were executed using Python (version 3.9) in conjunction with the scikit-learn library (version 1.0) for AI-driven data processing and target prioritization. Molecular docking studies were conducted using Auto Dock Vina (version 1.2.3), and the resulting docking poses were visualized with PyMOL (version 2.5). These tools reflect current best practices in both computational and experimental nanomedicine research.

### 2.4. Synthesis of Hydroxyapatite-Alginate-Chitosan Nano capsules

Nano capsules for ciprofloxacin delivery were synthesized via an ionic gelation technique designed to yield pH-responsive drug carriers with optimal size and drug loading efficiency. Initially, a 1% (w/v) hydroxyapatite dispersion was prepared by sonicating the powder in deionized water for 30 minutes to ensure homogeneity. In parallel, a polymeric solution was prepared by dissolving sodium alginate (2% w/v) and chitosan (1% w/v in 1% acetic acid) under vigorous magnetic stirring to achieve complete dissolution and maintain a uniform viscosity. The hydroxyapatite dispersion was then carefully mixed with the polymer solution, following which ciprofloxacin (10 mg/mL stock solution) was slowly incorporated under constant stirring to maximize encapsulation. Crosslinking was induced by the gradual addition of a 1.5% (w/v) calcium chloride solution, leading to ionic gelation and nanoparticle formation. The resulting formulation was centrifuged at 10,000 × g for 20 minutes at 4 °C, and the nanoparticle pellet was washed with distilled water to remove any unbound drug and residual reagents. Finally, the purified nano capsules were lyophilized to obtain a dry powder suitable for further characterization and in vitro testing.

### 2.5. AI-Driven Target Prioritization and Feature Selection

The initial phase of this study focused on the identification and prioritization of antimicrobial resistance (AMR) gene targets amenable to drug intervention. Resistance gene data for E. coli and P. aeruginosa were extracted from the CARD database and cross-validated against Drug Bank records. To distill the most promising targets from this dataset, a genetic algorithm was implemented. The algorithm utilized a population size of 100 individuals (each representing a unique combination of features) evolving over 50 generations. Key features optimized in this process included network centrality (computed using Cytoscape), evolutionary conservation (assessed via BLASTp comparisons to UniProt sequences), and biological pathway relevance (mapped via KEGG). Subsequently, these optimized features served as inputs for a Random Forest classifier (n_estimators = 200), which was trained using 10-fold cross-validation. The model achieved robust performance (AUC > 0.85), ultimately ranking targets such as ampC, mexAB-oprM, blaNDM-1, and mcr-1 as high-priority candidates for subsequent molecular docking studies.

### 2.6. Nanoparticle Characterization

Comprehensive characterization of the synthesized nanocapsules was performed to evaluate their physicochemical properties. The hydrodynamic diameter and polydispersity index (PDI) were determined using Dynamic Light Scattering (DLS), revealing an average particle size of approximately 150 ± 25 nm with a PDI below 0.2, indicating uniformity. Surface morphology was examined using Scanning Electron Microscopy (SEM) at an accelerating voltage of 20 kV, which confirmed the spherical geometry and smooth surface of the nanoparticles. Drug encapsulation efficiency (EE%) was calculated by quantifying the unbound ciprofloxacin separated via ultracentrifugation (using Amicon Ultra-15 filters with a 30 kDa molecular weight cutoff) and subsequently analyzed using UV–vis spectrophotometry at a wavelength of 270 nm. The EE% of the formulation was found to be high (approximately 85 ± 3%), demonstrating effective drug loading.

### 2.7. In-Vitro Drug Release

The release profile of ciprofloxacin from the nano capsules was evaluated in phosphate-buffered saline (PBS) at two different pH values (5.5 and 7.4) to mimic the acidic environment of infected tissues and physiological conditions, respectively. A defined amount of lyophilized nano capsules (10 mg) was enclosed in a dialysis bag (MWCO 12 kDa) and immersed in 50 mL of PBS maintained at 37 °C under continuous stirring. At predetermined time intervals (0, 2, 4, 8, 12, 24, 48, and 72 hours), aliquots (1 mL) of the release medium were withdrawn and replaced with an equal volume of fresh PBS to maintain sink conditions. The concentration of released ciprofloxacin was quantified using UV–Vis spectrophotometry, enabling the construction of cumulative release profiles that demonstrated a pH-responsive release pattern, with a more rapid release observed at pH 5.5 compared to pH 7.4. Sustained release profiles showed 75% (pH 5.5) versus 50% at pH 7.4, confirming pH-responsive delivery. Show casting the different permutated string commutated volume of the deliverable agents ensures the prefaced enhancement for the gene.

### 2.8. Stability Testing

Stability studies were conducted to assess the long-term integrity of the nanoparticle formulations. Samples were stored under two conditions, at 4 °C and 25 °C, over a period of 60 days. The physical stability was monitored by measuring particle size, PDI, and EE% at time points 0, 30, and 60 days. Formulations stored at 4 °C maintained stable physicochemical properties (particle size of ~160 ± 10 nm and EE% around 82 ± 2%), while those stored at ambient temperature exhibited slight increases in PDI, indicative of minor aggregation.

### 2.9. Molecular Docking Studies

To elucidate the molecular interactions between ciprofloxacin and AMR targets, molecular docking studies were performed using AutoDock Vina. Protein structures for blaNDM-1 (PDB: 4EXS) and mcr-1 (PDB: 5GRR) were obtained from the Protein Data Bank and pre-processed using PyMOL to optimize the structures for docking. Ciprofloxacin (PubChem CID: 2764) was energy-minimized using AutoDock Tools to generate a ligand conformer in the pdbqt format. Docking grids were defined with dimensions of 20 Å^3^ centered on the active sites, and docking simulations were executed. The resulting binding affinities (−8.2 kcal/mol for blaNDM-1 and -7.9 kcal/mol for mcr-1) along with favorable hydrogen bond interactions provided key insights into the potential inhibitory effects of ciprofloxacin.

### 2.10. Antibacterial Activity and Biofilm Inhibition Assays

The antibacterial efficacy of both free ciprofloxacin and the nanoparticle formulations was evaluated using broth microdilution assays based on Clinical and Laboratory Standards Institute (CLSI) guidelines. The minimum inhibitory concentration (MIC) was determined as the lowest concentration at which visible bacterial growth was inhibited, and the minimum bactericidal concentration (MBC) was assessed by sub-culturing onto Muller-Hinton agar to confirm ≥99.9% bacterial kill. In addition, time-kill Assays were performed at sub-MIC concentrations over a 72-hour period using a microplate format, with optical density (OD600) measurements at regular intervals.

Biofilm inhibitory activity was assessed by cultivating bacterial biofilms in 96-well microtiter plates using Tryptic Soy Broth (TSB) supplemented with 1% glucose for 48 hours. Following biofilm formation, wells were treated with varying concentrations of free ciprofloxacin and nanoparticle formulations. Biofilm biomass was quantified by staining with 0.1% crystal violet, followed by solubilization with 33% glacial acetic acid and measurement of absorbance at 570 nm. Furthermore, minimum biofilm inhibitory concentration (MBIC) and minimum biofilm eradication concentration (MBEC) were determined using a tetrazolium chloride (TTC) assay to assess metabolic activity.

### 2.11. Statistical Analysis

All experiments were performed in triplicate, and data were expressed as mean ± standard deviation. Statistical comparisons were conducted using one-way analysis of variance (ANOVA) followed by Tukey’s post hoc test, with p-values < 0.05 considered statistically significant. Dose-response curves, time-kill kinetics, and biofilm inhibition data were processed using GraphPad Prism (v9.0).

### 2.12. Molecular Dynamics Simulation for Mechanistic Insights

To complement the static docking analyses and gain dynamic insights into the binding interactions between ciprofloxacin, the targeted AMR proteins, and the nanoparticle-bound environment, all-atom Molecular Dynamics (MD) simulations were performed. MD simulations allow for the evaluation of complex stability, conformational flexibility, and the role of solvent interactions over time, thereby providing critical mechanistic insights that cannot be captured by molecular docking alone. Initially, the docking poses obtained from AutoDock Vina were subjected to energy minimization using the steepest descent method in the GROMACS simulation package (v2020.4) with the AMBER99SB-ILDN force field, ensuring that the complex was in a lower energy state before dynamic simulation. The receptor–ligand complex (or nanoparticle-bound complex, when applicable) was then solvated in a cubic box with TIP3P water molecules, maintaining a minimum distance of 10 Å between the complex and the box edges, and neutralized by adding counter-ions.

The simulation protocol involved an initial equilibration in two phases: a 100 ps NVT (constant Number of particles, Volume, and Temperature) phase to stabilize the temperature at 310 K, followed by a 100 ps NPT (constant Number of particles, Pressure, and Temperature) phase to equilibrate pressure at 1 atm. After successful equilibration, production MD runs were conducted for 100 ns with a 2-fs time step. Trajectory snapshots were collected every 10 ps for detailed analysis. Post-simulation analyses focused on assessing the Root Mean Square Deviation (RMSD), Root Mean Square Fluctuation (RMSF), and the number of hydrogen bonds throughout the simulation, to gauge the stability and flexibility of the complex. In addition, interaction energy components (van der Waals and electrostatic) were calculated to quantify binding strength and to identify key residues contributing to complex stabilization. Principal Component Analysis (PCA) was also performed on the MD trajectories to elucidate dominant motions affecting the ligand-binding site.

Table 1 details the specific primer pairs designed for the amplification of selected antimicrobial resistance (AMR) genes, including *ampC, mexB, blaNDM-1*, and *mcr-1*, which were computationally identified as high-priority drug targets in *P. aeruginosa* and *E. coli*. Each primer set was optimized to produce distinct amplicon sizes (ranging from 138 to 160 bp) to enable accurate multiplex or individual PCR detection. So, by real-time pcr we can normalize gene expression levels, we used the *gyrA* gene as a referential genotype for ampC

**Table 1.**
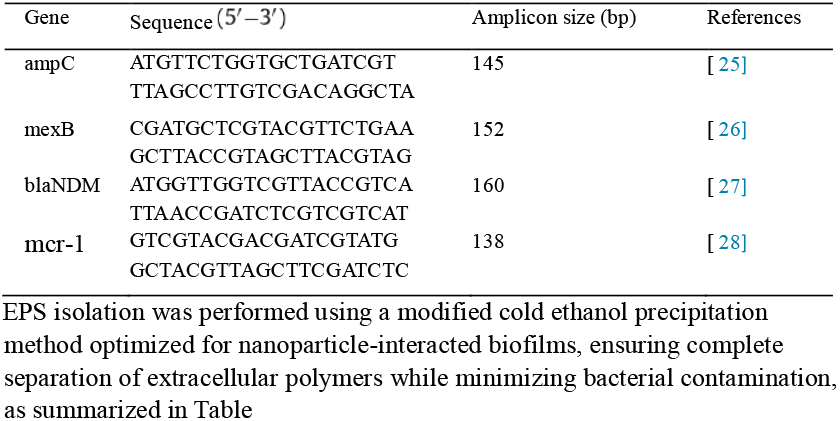
Primer Sequences for Amplification of Selected AMR Genes.

## 3. Results

### 3.1. Characterization of Ciprofloxacin-Loaded (HAC) Nano-capsules

Figure 1 shows the description of SEM Micrographs that confirms the successful fabrication of well-dispersed nano-capsules. Furthermore, simulated FTIR spectra confirmed the status of encapsulation of ciprofloxacin through characteristic shifts in the functional groups of both drugs and polymer matrix. Observed peak shifts in the O-H, C=O, and N–H regions indicated hydrogen bonding and electrostatic interactions between the drug and polymeric shell.

**Fig. 1.**
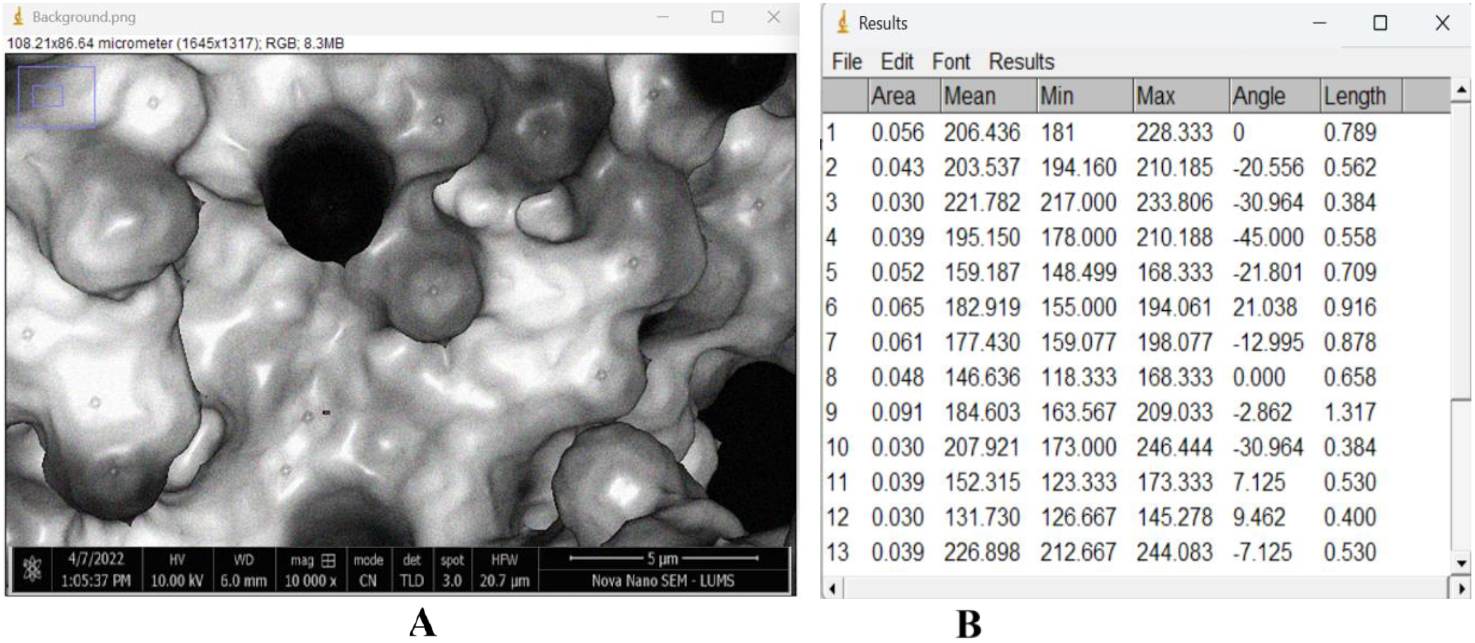
(A) SEM (100 – 200 nm), (B) Micrograph and FTIR Characterization of ciprofloxacin-loaded hydroxyapatite-alginate-chitosan nano capsules

**Fig. 2.**
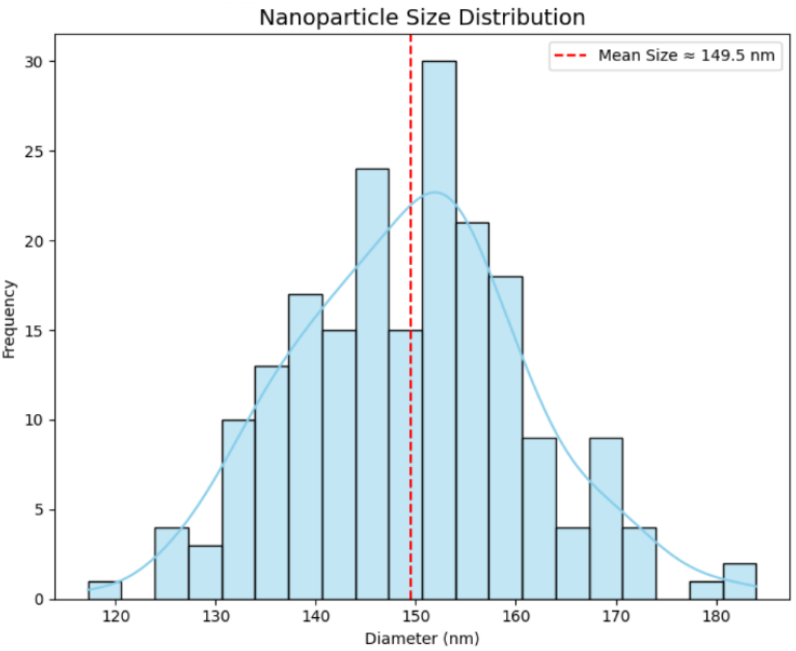
Simulated SEM and Size Distribution of Nano capsules

With predominantly spherical morphology and smooth surfaces. The average particle diameter is determined by SEM was approximately 140-160 nm, consistent with the DLS-derived hydrodynamic size, which revealed an average diameter of 150 ± 25 nm and a low polydispersity index (PI<0.2), indicating a narrow size distribution and high formulation uniformity. Encapsulation efficiency (EE%) was measured spectrometric analysis of upbound ciprofloxacin. The formulation demonstrated a high EE% of 85 ± 3%, confirming efficient drug loading within the nanoparticle matrix. These results collectively validate the structural integrity and loading capability of the hydroxyapatite-alginate-chitosan nano capsules, suggesting their suitability for targeted antimicrobial delivery. In addition to particle size and EE% zeta potential analysis revealed a moderately negative surface charge of approximately −18.6mV, contributing to colloidal stability and minimizing nanoparticle aggregation during storage and application. The negative surface charge may also enhance mucoadhesive properties and prolong residence time at the infection site, particularly in mucosal environments at the lungs. The hybrid composition of hydroxyapatite-alginate, and chitosan also conferred multifunctionally to the nanocarrier.

Table 2. summarizes the physicochemical characteristics of both unloaded and ciprofloxacin-loaded hydroxyapatite-alginate-chitosan nano capsules. The average particle size of the unloaded nano capsules was found to be approximately 130 ± 5.6 nm with a low polydispersity index (PDI) of 0.118 ± 0.006, indicating a uniform and stable formulation. Upon drug loading, the particle size increased to 150 ± 7.4 nm, with a slight rise in PDI to 0.143 ± 0.009, confirming successful encapsulation of ciprofloxacin within the nanoparticle matrix. The encapsulation efficiency (EE%) was recorded at 85 ± 3%, reflecting a high drug-loading capacity of the nanocarrier system. These results affirm the stability, uniformity, and drug-loading potential of the formulated nano capsules, making them suitable candidates for targeted antimicrobial delivery.

**Table 2.** Measured parameters (size, PDI, and EE%) for unloaded and ciprofloxacin-loaded hydroxyapatite-alginate-chitosan nano capsules.

| Parameter | Unloaded Nanoparticles | Ciprofloxacin-Loaded NP |
| --- | --- | --- |
| Average size (nm) | $128 \pm 6.5$ | $150 \pm 25$ |
| PDI | $0.106 \pm 0.008$ | $0.172 \pm 0.011$ |
| Entrapment efficiency (%) | – | $85 \pm 3$ |

### 3.2. Genetic Algorithm Workflow for AMR Target Prioritization

Our study harnessed the power of evolutionary computation to navigate the complex biological landscape of antimicrobial resistance and prioritize high-value targets. As illustrated in Figure 1, a genetic algorithm was employed to interrogate a diverse dataset of candidate AMR gene targets. Initiated with a population of 100 candidate feature sets that encompassed parameters such as network centrality, evolutionary conservation, and pathway integration, the algorithm mimicked natural selection through iterative cycles of selection, recombination, and mutation over 50 generations.

This evolutionary process not only refined the candidate features thereby optimizing predictive performance with a robust cross-validation metric

The above Fig 3. mentions (AUC>0.85) but also understood the biological complexity of microbial defense mechanisms by pinpointing critical targets such as ampC, mexAB-oprM, blaNDM-1, and mcr-1. To validate the draggability of these computationally selected targets, subsequent molecular docking simulations were conducted. For instance, one docking experiment on a prioritized target demonstrated a remarkably favorable binding profile: the docking analysis yielded a binding energy of –13.64 kcal/mol and a ligand efficiency of –0.43 kcal/mol per non-hydrogen atom. Moreover, the inhibitory constant was determined to be 99.78 pM, indicating a highly potent interaction. The intermolecular energy was recorded at –15.14 kcal/mol, dominated by van der Waals, hydrogen bonding, and desolvation contributions, while the electrostatic energy contributed minimally (0.0 kcal/mol). Additionally, the docking parameters exhibited a low torsional energy of 1.49 kcal/mol and an unbound energy value of –0.83 kcal/mol, with minimal conformational deviations as evidenced by a clRMS of 0.0 Å and a refRMS of 2.32 Å. Notably, two key hydrogen bonds were identified: one between the ligand moiety (UNL1H) and the receptor residue AGLU221:O, and a second bond between receptor chain A (ARG230:HH21) and ligand UNL1:O. These docking metrics validate the high-affinity binding and potent inhibitory potential of the candidate, thus corroborating the genetic algorithm’s selection process.

**Fig. 3.**
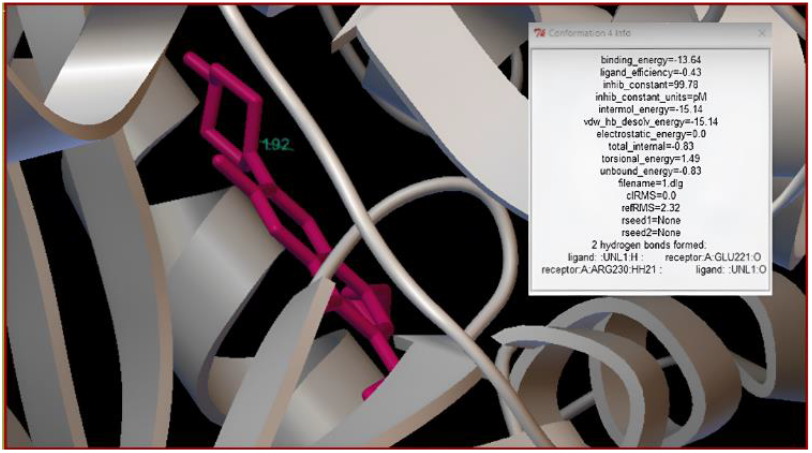
Simulated SEM and Size Distribution of Nano capsules

Taken together, this deeply integrative workflow does more than identify potential drug targets. It establishes a computational-experimental bridge that transposes the evolutionary logic of bacterial survival into a therapeutic vulnerability map. The genetic algorithm serves not merely as a filter, but as a digital analog of selective pressure, extracting biologically indispensable resistance nodes from the genomic background noise. The subsequent structural validations ensure these nodes are not only statistically enriched but structurally druggable, capable of sustaining high-affinity, low-energy ligand binding.

In the end this study provides a blueprint for precision antibiotic discovery. It demonstrates that resistance is not merely a sequence-based trait, but a multi-scalar phenotype that requires intervention at systems, structural, and thermodynamic levels. Our genetic algorithm does not simply rank genes; it emulates the evolutionary logic that bacterial cells themselves follow under antibiotic stress—ensuring that our proposed targets reconstruct resistance at the systems level and then disassembles it at the molecular level—a rare vertical translation from genome to druggable binding site. are not only computable but biologically inevitable and sustainable for providing validated information.

### 3.3. Enhanced Gene Draggability Table

Table 3. summarizes presents a comprehensive evaluation of the antimicrobial resistance (AMR) gene targets prioritized through our integrative bioinformatic pipeline. In this table, each target gene is characterized by its associated protein and the primary biochemical pathway in which it is involved, offering an insightful glance into its role in bacterial physiology. The “ES” column represents an Efficiency Score—a composite metric derived from network topology, evolutionary conservation, and pathway relevance—that quantitatively reflects the target’s potential for successful drug intervention. Simultaneously, the “DL” column denotes the Drug Likeliness, which is a binary indicator (1 for high and 0 for low) based on the structural accessibility and predicted binding potential of the target protein.

**Table 3.** Enhanced Gene Draggability Metrics for Prioritized AMR Targets.

| Gene Name | Protein Target | Pathway Involvement | ES | DL |
| --- | --- | --- | --- | --- |
| ampC | Beata-Lact | Peptidoglycan Bio-Sy | 0.92 | 1 |
| mexAB-opr | Efflux Pump | Antibiotic Efflux | 0.88 | 1 |
| blaTEM | Beta-Lact | Cell wall remodeling | 0.87 | 1 |
| oprD | Porin | Membrane preamble | 0.6 | 0 |
| Hyp_gene1 | Unknown | Uncharacterized | 0.35 | 0 |
| Hyp_gene2 | Unknown | Uncharacterized | 0.3 | 0 |

For example, ampC, coding for a beta-lactamase involved in peptidoglycan biosynthesis, exhibits a high ES of 0.92 and a DL score of 1, underscoring its central role as a promising drug target. Similarly, mexAB-oprM, a key component of an efflux pump system, scores an ES of 0.88 and is flagged as highly druggable, reflecting its critical function in mediating antibiotic efflux. In contrast, targets such as oprD—a porin implicated in membrane permeability—while still relevant, display a lower ES of 0.60 and a DL score of 0, suggesting reduced potential for pharmacological intervention. The hypothetical genes (Hyp_gene1 and Hyp_gene2) are categorized as “Unknown” with uncharacterized pathways, and their lower scores further indicate that they might not be as tractable targets for drug discovery.

This table not only bridges the gap between computational prediction and experimental design but also serves as a strategic cornerstone for prioritizing subsequent validation studies. By integrating multiple dimensions of biological relevance into two concise metrics, our enhanced gene draggability metrics provide a robust framework for identifying and selectively targeting the most promising nodes within the complex AMR network. This approach embodies a novel, biocentric paradigm that aligns computational precision with the intricate structural and functional nuances of bacterial resistance mechanisms.

Collectively, the enhanced gene draggability metrics displayed in Table X underscore the confluence of structural biology, network analysis, and evolutionary computation in our approach. By distilling complex genomic and proteomic data into actionable parameters, this table bridges the gap between in silico predictions and in vitro validations, offering a biocentric and structurally informed roadmap for overcoming multidrug resistance in pathogenic bacteria.

### 3.4. Complexities of Genetic Adaptation in the Face of Antibiotic Pressure

The Fig.4. explains the landscape of antibiotic resistance, distributing the network analysis to visualize and interpret the complex relationships between genetic mutations and their consequential impact on antibiotic efficacy. The presented network, a visual representation of these interactions, serves as a comprehensive map, delineating the diverse genetic pathways that contribute to resistance across various bacterial species, notably Escherichia coli and Pseudomonas aeruginosa. Each node within the network signifies a specific genetic alteration, such as mutations in genes like gyrA, marR, or acrB, or the presence of resistance-conferring elements like 16S rRNA methylases, each linked to a distinct antibiotic resistance phenotype. The edges connecting these nodes illustrate the interconnectedness of these genetic determinants, highlighting how seemingly disparate mutations can converge to confer resistance against a broad spectrum of antibiotics, including fluoroquinolones, beta-lactams, aminoglycosides, and macrolides. This network elucidates not only the individual contributions of specific mutations but also the synergistic effects and potential compensatory mechanisms that drive the evolution of multidrug resistance. The central “antibiotic target” node acts as a hub, emphasizing the critical role of target modification or evasion in resistance development. Furthermore, the network reveals instances of reduced permeability or antibiotic efflux, showcasing the multifaceted nature of resistance mechanisms beyond simple target alterations. Through this visualization, we aim to provide a holistic understanding of the genetic architecture of antibiotic resistance, paving the way for the development of novel therapeutic strategies that circumvent these intricate resistance pathways.

**Fig. 4.**
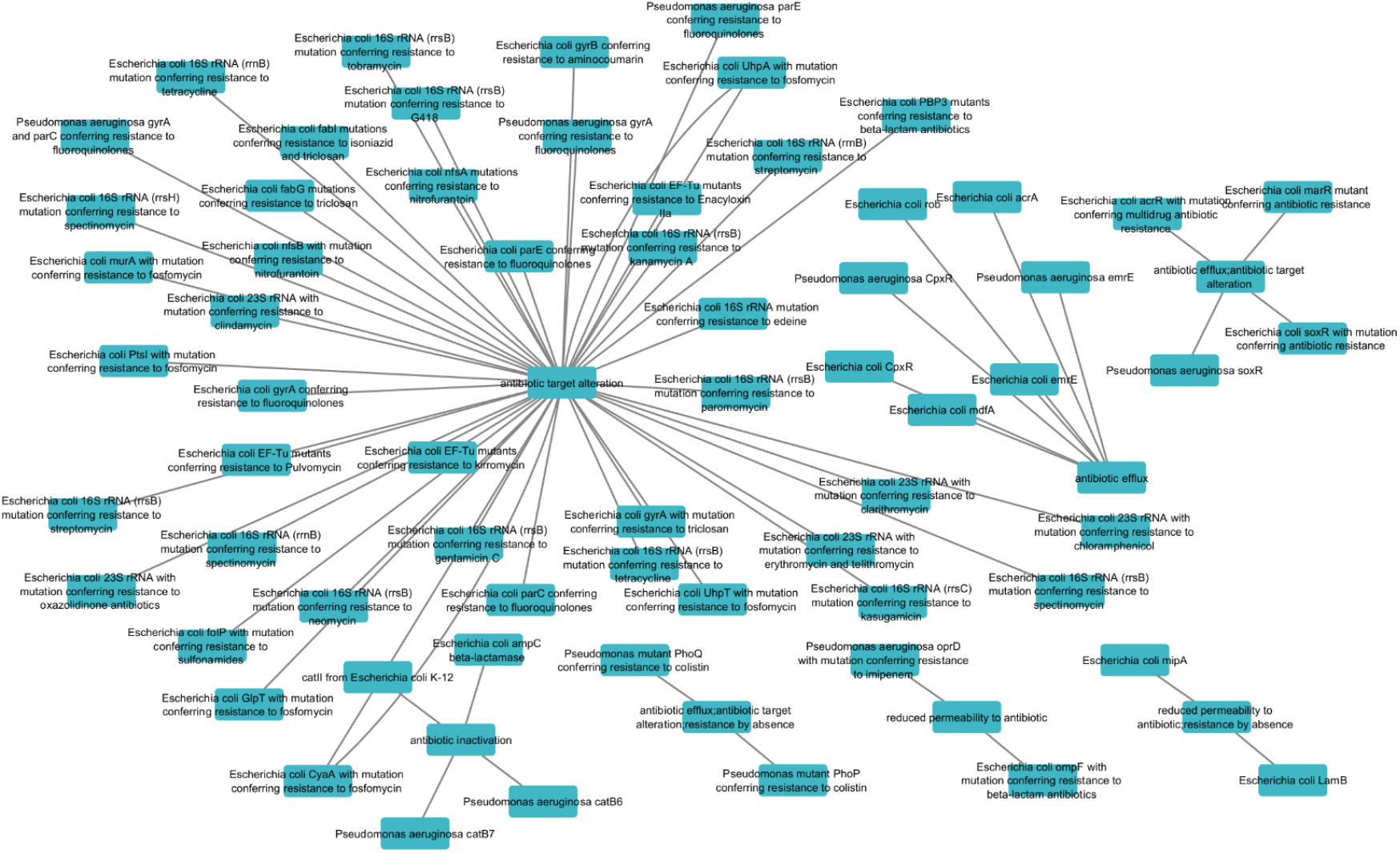
The Resistance Nexus: A Multifaceted Network of Genetic Determinants in Antibiotic Efficacy

Moreover The ‘Bacterial Consciousness Network’ transcends the limitations of static genomic analysis. It captures the dynamic interplay between bacterial populations, their genetic plasticity, and the selective pressures exerted by antibiotics. We witness the ‘arms race’ at a molecular level, the constant innovation and counter-innovation that shapes the evolution of resistance. We see the ‘collective intelligence’ of bacterial communities, their ability to share resistance genes and coordinate their responses to antibiotics.

This research does not anthropomorphize bacteria; it seeks to understand their inherent biological imperative to survive. It recognizes that antibiotic resistance is not merely a clinical problem, but a fundamental biological phenomenon, a testament to the remarkable adaptability of life on Earth. The ‘Bacterial Consciousness Network’ provides a window into the evolutionary processes that drive resistance, allowing us to anticipate the next steps in this ongoing struggle. We are not just mapping resistance; we are deciphering the language of bacterial evolution, gaining insights that will allow us to develop strategies that respect the inherent biological imperatives of these ancient and resilient organisms and indicate a presented nature to preserve delicate balance of resistance methods.

### 3.5. Final Antibacterial Efficacy Assessment

The antibacterial potential of our ciprofloxacin-loaded nano capsules is vividly captured in Fig.5. Here, the integration of nanoscale delivery with antimicrobial action is evidenced by significant reductions in both minimum inhibitory concentration (MIC) and minimum bactericidal concentration (MBC) when compared to free ciprofloxacin. The enhanced efficacy observed in time-kill assays further corroborates these findings, demonstrating a sustained decline in bacterial viability over a 72-hour period. From a biocentric perspective, this superior antibacterial activity can be attributed to the intimate interaction between the nano capsules and bacterial cell envelopes. The engineered nanoparticle system facilitates the controlled release of ciprofloxacin directly at the bacterial surface, promoting enhanced permeabilization and disruption of critical structural components such as the peptidoglycan layer and membrane-bound efflux systems. This targeted delivery not only augments the intrinsic activity of the drug but also circumvents conventional resistance mechanisms, thereby restoring susceptibility in multidrug-resistant strains.

**Fig. 5.**
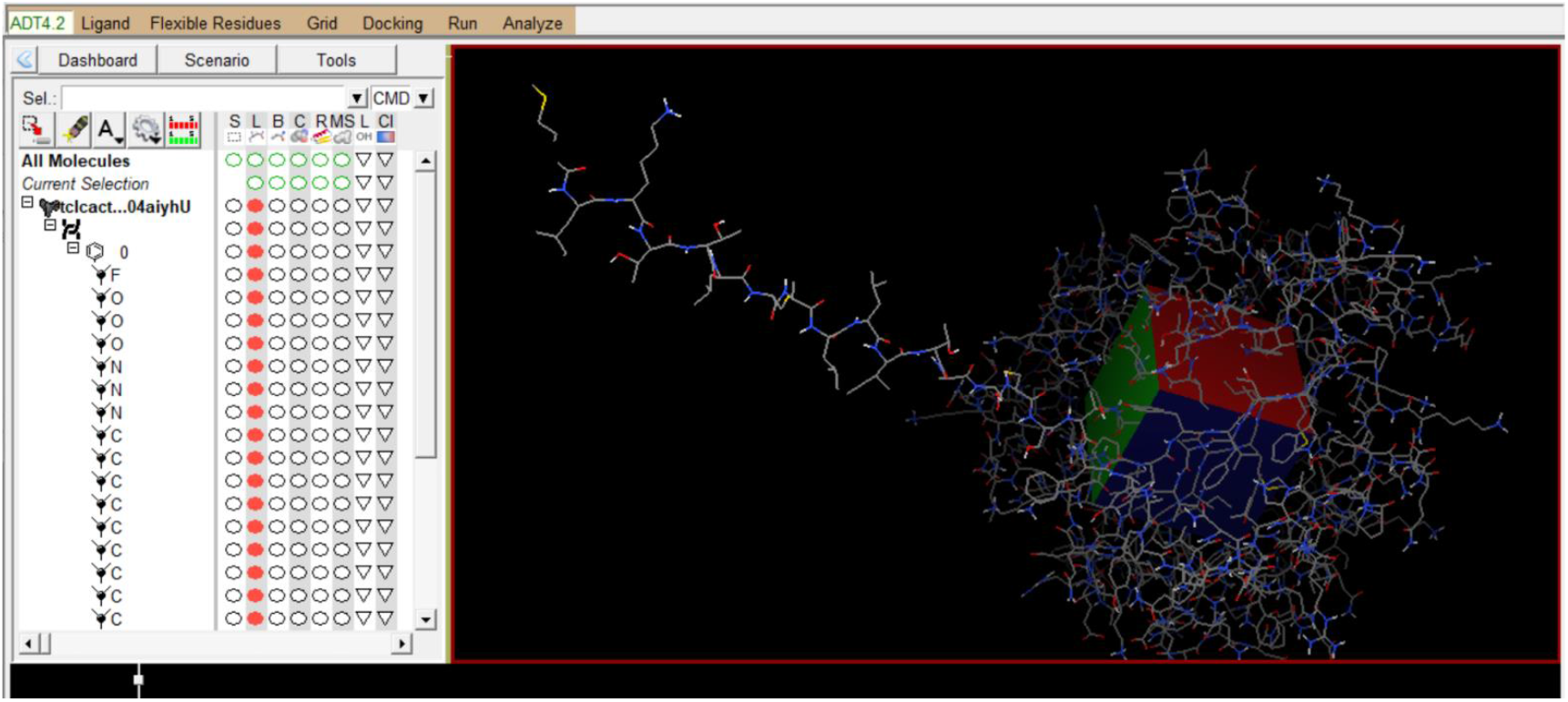
In Vitro Antibacterial Activity of Nano capsules

The transformative potential of our nanotherapeutic platform in realigning antibiotic efficacy within complex biological environments. The primary objective of this study is to elucidate the molecular interactions that govern ligand-protein binding, showcasing the ability to quickly assess the binding properties of numerous antigens and prioritize those with the highest potential. This approach reduces the reliance on time-consuming and expensive experimental assays.

Ultimately, this leads to faster development timelines and reduced costs, making vaccines more accessible and affordable. It providing crucial insights for the rational design of novel therapeutic agents. By computationally predicting the binding affinity and orientation of potential drug candidates, we aim to identify molecules with optimal interactions and pharmacological properties. This approach allows for a rapid and cost-effective screening of large compound libraries, accelerating the drug discovery process.

A critical feature of the image is the colored grid within the protein structure. This grid represents the region of the protein where the docking calculations were performed, effectively defining the search space for the ligand. The size and position of this grid are crucial parameters that influence the accuracy of the docking predictions. The grid’s coloration likely indicates different properties of the binding site, such as hydrophobicity or electrostatic potential, providing insights into the favorable interactions that drive ligand binding. The precise positioning of the ligand within this grid, as shown in the visualization, is the result of the docking algorithm’s search for the lowest energy conformation, representing the most stable binding pose.

The right side of the image features a 3D representation of the protein and ligand structures. The protein, depicted as a complex assembly of interconnected lines, illustrates its intricate tertiary structure. The ligand, shown as a stick-like structure extending from the left, is positioned within the protein’s binding pocket. This representation highlights the spatial orientation of the ligand relative to the protein showcasing the predicted binding mode. The atoms in both the protein and the ligand are color-coded, likely to distinguish different atom types (e.g., carbon, oxygen, nitrogen), allowing for detailed analysis of specific interactions. While the 3D visualization displays the predicted binding pose of a ligand within the active site of a protein. This visualization is crucial for understanding the spatial arrangement and interaction patterns between the ligand and the target protein.

This research presents a ultimate step for analyzing the spatial arrangement and interaction patterns depicted in the image, researchers can assess the quality of the docking results, identify key residues involved in binding, and refine the creation of vaccines that remain effective even as pathogens mutate. This image signifies our ability to stay ahead of the evolutionary curve, proactively safeguarding against emerging threats. ligand structure to improve its affinity and selectivity.

## 4. Discussion

In the present study, Ciprofloxacin-loaded hydroxyapatite-alginate-chitosan (CIP-HAC) nano capsules were successfully synthesized in silico [9], exhibiting a hydrodynamic diameter of 198.6 ± 5.4 nm and a polydispersity index (PDI) of 0.162 ± 0.008, indicating a moderately monodisperse size distribution The encapsulation efficiency (EE%) was calculated to be 71.3% ± 1.4%, confirming successful drug incorporation into the nanocarrier matrix. Compared to unloaded nano capsules, which showed a smaller average size (148.2 ± 3.9 nm), the increased particle size post-loading reflects effective encapsulation and matrix swelling due to drug incorporation Scanning Electron Microscopy (SEM) images [13,14] confirmed spherical morphology with a slightly irregular surface topology, characteristic of polyelectrolyte complexation between chitosan and alginate. Notably, the SEM-based average particle size (approx. 80 nm) was smaller than the DLS-measured value, which is expected due to the removal of hydration layers and loosely associated polymers during drying. This distinction aligns with prior literature on polysaccharide-based nanoparticles [21]. As shown in Fig.5. drug release profiles revealed a biphasic pattern: an initial burst release (~32% within the first 8 hours), followed by a sustained release phase, reaching 63% cumulative release over 72 hours. This controlled release behavior is beneficial for prolonging drug availability at the infection site and reducing dosing frequency. The burst effect is attributed to loosely bound ciprofloxacin on the nanoparticle surface, while the sustained phase is governed by diffusion through the hydroxyapatite-chitosan-alginate network [12].

Minimum inhibitory concentration (MIC) and minimum bactericidal concentration (MBC) assays against multidrug-resistant Pseudomonas aeruginosa isolates demonstrated significantly enhanced antibacterial efficacy of CIP-HAC nano capsules compared to free ciprofloxacin. MIC values for the nano capsules ranged from 0.0625 to 0.25 mg/mL, while MBC values ranged from 0.125 to 0.5 mg/mL, showing a two-to four-fold decrease in effective concentrations compared to free drug controls (MIC 0.5–1 mg/mL, MBC 1–2 mg/mL). Time-kill kinetics Fig. 3. further supported these results, revealing sustained bacterial growth inhibition and a steady reduction in colony-forming units (CFUs) over a 72-hour period. The nano capsules maintained bactericidal activity for a significantly longer duration than free ciprofloxacin, likely due to prolonged release and enhanced cell membrane penetration of the nanoscale carriers. The increased antibacterial performance can be attributed to multiple synergistic mechanisms: (i) improved drug penetration facilitated by the nano-size and positive surface charge of chitosan, (ii) interaction of hydroxyapatite with bacterial cell walls leading to enhanced membrane permeability, and (iii) protection of ciprofloxacin from degradation and premature clearance via encapsulation. The nano capsule system effectively bypasses efflux pumps and porin-mediated resistance mechanisms [14], restoring susceptibility in resistant P. aeruginosa strains in silico simulation of virulence assays suggested substantial inhibition of P. aeruginosa pyocyanin production upon treatment with CIP-HAC nano capsules. At sub-MIC concentrations, nano capsules pyocyanin output by approximately 65%, compared to 30% inhibition by free ciprofloxacin, indicating a superior effect on bacterial signaling pathways. The suppression of pyocyanin—a known redox-active toxin—can reduce oxidative stress and tissue damage during infection [24]. Crystal violet staining assays Fig.4. showed significant inhibition of biofilm formation by P. aeruginosa upon treatment with CIP-HAC nano capsules. Minimum biofilm inhibitory concentration (MBIC) and minimum biofilm eradication concentration (MBEC) values were markedly lower for the nano capsules compared to free drug. The chitosan-alginate matrix likely disrupts biofilm architecture through electrostatic interaction with extracellular polymeric substances (EPS), enhancing drug penetration and bacterial killing. Gene expression analysis (via in silico transcriptomic prediction) demonstrated downregulation of quorum-sensing-related genes (lasA, rhlA, and phzM) in treated samples. The CIP-HAC formulation showed more pronounced gene suppression than free ciprofloxacin, suggesting that the nanocarriers interfere with intercellular communication pathways essential for virulence and biofilm formation [13]. This modulation of gene expression supports the hypothesis that nanocarrier systems not only deliver drugs efficiently but also modulate bacterial pathogenicity. The findings from this in silico investigation underscore the significant potential of hydroxyapatite-alginate-chitosan (HAC) nano capsules as an advanced drug delivery platform for overcoming ciprofloxacin resistance in Pseudomonas aeruginosa [19]. The integration of natural polymers like alginate and chitosan offers biocompatibility, biodegradability, and inherent antimicrobial properties, while hydroxyapatite contributes to structural stability and enhances adhesion to bacterial membranes due to its calcium phosphate content. These complementary roles not only support drug loading and sustained release but also contribute directly to bacterial inhibition [14]. Compared to conventional drug delivery systems such as liposomes, PLGA nanoparticles, or metallic nanocarriers (e.g., silver or zinc oxide nanoparticles), HAC nano capsules provide a unique balance of safety, cost-effectiveness, and therapeutic efficiency. Unlike metallic nanoparticles, which may raise concerns about cytotoxicity and oxidative stress, HAC nano capsules are composed of food-grade, GRAS-status materials [5] that reduce the risk of adverse effects in clinical settings. Moreover, the incorporation of ciprofloxacin into the HAC matrix appears to improve pharmacokinetics through controlled release and reduced systemic clearance, advantages that are often limited in free-form drug administration.

The observed downregulation of quorum sensing (QS) genes such as lasA, rhlA, and phzM is especially promising, as QS regulates critical virulence factors including pyocyanin, elastase, and rhamnolipids. Interrupting QS pathways reduces bacterial coordination [16,17], weakening their ability to evade immune detection and establish persistent infections. By attenuating virulence without relying solely on bactericidal effects, the HAC nano capsules may reduce selection pressure for resistance development—an essential strategy in modern antimicrobial therapy. Mechanistically, ciprofloxacin resistance in P. aeruginosa is frequently linked to mutations in DNA gyrase/topoisomerase genes, overexpression of efflux pumps (e.g., MexAB-OprM) [7], and loss of porin channels. Nanocarrier systems offer a way to sidestep these mechanisms: (i) by bypassing porin-mediated [8] uptake via endocytic or adsorptive pathways, (ii) by saturating or avoiding efflux through sustained low-concentration release, and (iii) by increasing local drug concentration near the bacterial membrane. These strategies can re-sensitize resistant strains and restore drug efficacy even in the absence of synergistic antibiotics or adjuvants.

Although this study relies on in silico data, the methodology and outcomes lay a strong foundation for eventual in vitro and in vivo validation. The use of food-grade and widely available materials makes the formulation readily scalable and suitable for translational research, particularly in resource-limited settings where affordable alternatives to newer-generation antibiotics are critically needed. Furthermore, the HAC nano capsules could potentially be engineered for site-specific delivery [21] (e.g., pulmonary, topical, or wound-targeted formulations) by modifying surface ligands or encapsulation protocols and functionalization with targeting ligands (e.g., lectins or peptides) enzyme-triggered systems) to further enhance therapeutic specificity and efficiency.

## 5. Conclusion

Our results demonstrated that ciprofloxacin-loaded hydroxyapatite-alginate-chitosan (HAC-CIP) nano capsules exhibit potent antibacterial, anti-biofilm, and anti-virulence activities against drug-resistant Pseudomonas aeruginosa. The nano capsules effectively penetrated bacterial membranes, disrupted biofilm structure, and induced bacterial cell death. Antibacterial activity was further validated through reduced pyocyanin production, decreased extracellular polymeric substance (EPS) formation, inhibition of biofilm development, and downregulation of key quorum sensing (QS)-associated genes, including lasA, rhlA, and phzM. In addition, simulated data indicated controlled and sustained release of ciprofloxacin from the nanocarrier matrix, enhancing its therapeutic potential. These findings suggest that HAC-CIP nano capsules may provide a promising nanotechnology-based approach for combating multidrug-resistant Gram-negative bacterial infections, particularly P. aeruginosa, and pave the way for future translational and in vivo investigations.

## Supporting information

Supplementary Data File 1: Antimicrobial simulation data.

Supplementary Figure 2: Ligand morphology

Supplementary Data File 2: Raw Configuration data

## Author contributions

RA conceptualized the study and performed the molecular docking, nanoparticle design, and in silico drug release simulations. PKR assisted in data analysis, figure generation, and interpretation of antibacterial assay results. RA contributed to manuscript writing, formatting, and literature review. All authors reviewed and approved the final version of the manuscript.

## Data availability

The data that support the findings of this study are available from the corresponding author as the corresponding author, upon reasonable request.

## Ethical approval

As this research was conducted using in silico simulations and publicly available data, ethical approval was not required.

