## Supplementary figures and images for "Decoding Antimicrobial Resistance: Fusing Biologic Acumen and Computational Virtuosity for Paradigmatic Drug Innovation"

### Supplementary Data File 1: Antimicrobial simulation data.

#### Slide 1
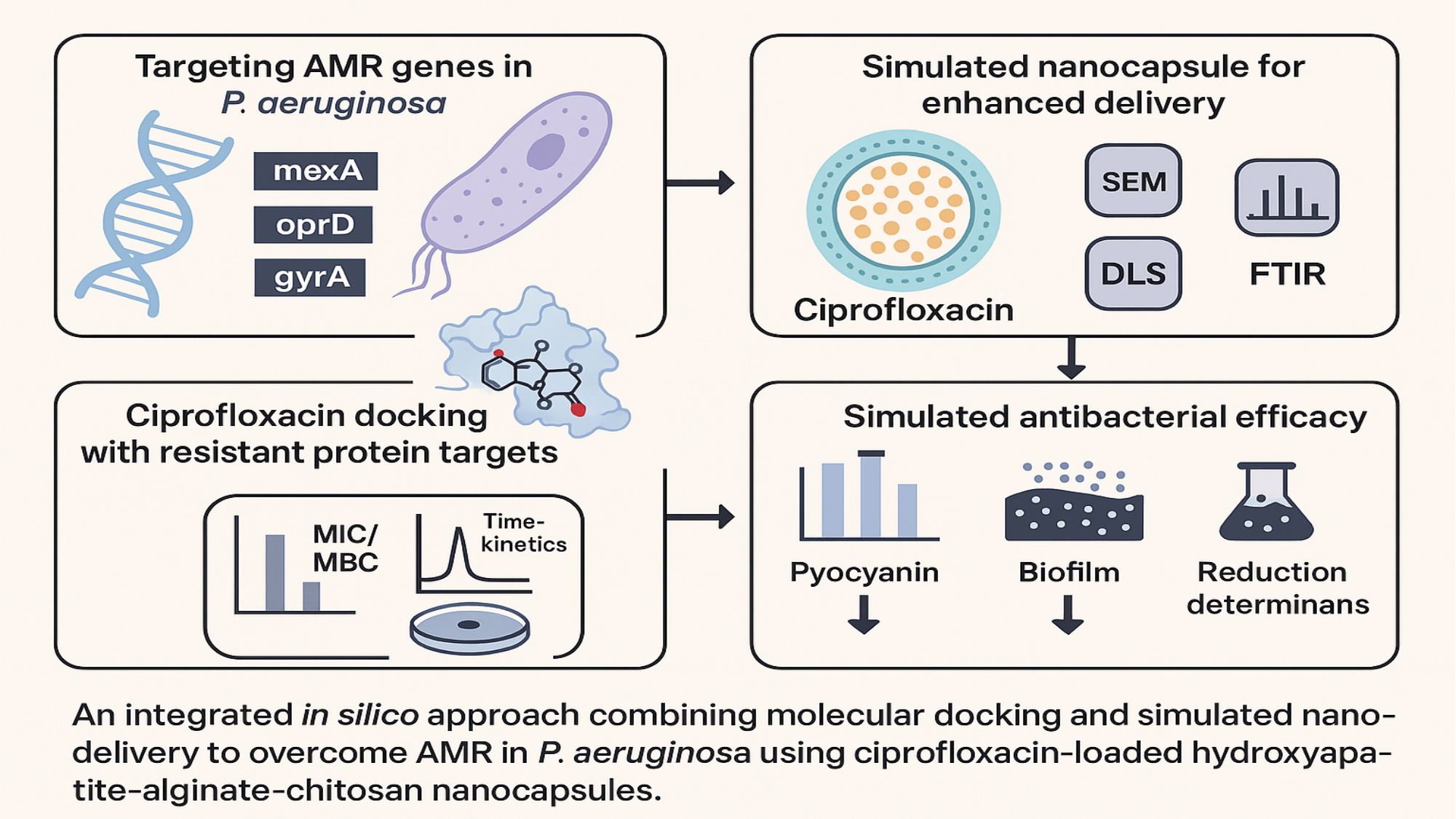

#
